# Ferrosome Organelles Compartmentalize a Redox-Active Ferrous Phosphate Biomineral

**DOI:** 10.64898/2026.06.16.732667

**Authors:** Kewei Zhao, Kristina M. Ferrara, Yixuan Liu, Macon J. Abernathy, Ritimukta Sarangi, Hualiang Pi

**Author notes:** Correspondence should be addressed to Ritimukta Sarangi or Hualiang Pi.;. These authors contributed equally (Kewei Zhao, Kristina M. Ferrara).

## Abstract

Ferrosomes are recently discovered lipid-bound bacterial organelles that store iron as iron-phosphate biominerals, yet the chemical nature and physiological consequences of ferrosome-stored iron remain poorly understood. Here, we combined X-ray absorption spectroscopy (XAS), electron microscopy, inductively coupled plasma mass spectrometry (ICP-MS), and physiological analyses to characterize ferrosome iron in *Clostridioides difficile*. We initially characterized isolated ferrosomes from a *fur*::CT strain that overproduces these organelles. XAS identified an amorphous iron-phosphate biomineral containing mixed Fe(II)/Fe(III), consistent with partial oxidation during aerobic isolation. In contrast, whole-cell XAS of anaerobically maintained *fur*::CT cells revealed predominantly ferrous iron with local Fe–O–P coordination features resembling those of vivianite. Similar spectral features were observed in wild-type cells under conditions that promote ferrosome abundance. Additionally, deletion of the *fez* operon abolished these characteristic spectral signatures, whereas complementation restored them. Upon air exposure, ferrosome-associated iron rapidly oxidized to a ferric phosphate-like state, demonstrating the oxygen sensitivity of this mineral phase. Physiological analyses further showed that ferrosome overproduction did not increase sensitivity to short-term oxygen exposure but increased susceptibility to high concentrations of H_2_O_2_ in a ferrosome-dependent manner. Together, these results establish ferrosomes as iron-storage organelles that sequester redox-active Fe(II) in a mineralized ferrous phosphate phase, thereby limiting its participation in cytosolic ROS chemistry and providing a mechanism for the safe storage of reactive iron.

**Significance Statement:** Iron is essential for life but can also damage cells because ferrous iron promotes oxidative chemistry. How cells store large amounts of ferrous iron while limiting its potential toxicity remains a fundamental biological question. Ferrosomes are recently discovered bacterial organelles that store iron as iron-phosphate biominerals, but the chemical nature and physiological consequences of ferrosome-associated iron remained unknown. Using Fe K-edge X-ray absorption spectroscopy, we show that ferrosomes in *Clostridioides difficile* contain a vivianite-like ferrous phosphate biomineral. Our data support a model in which ferrosome compartmentalization limits the participation of stored iron in cytosolic ROS chemistry. These findings reveal how bacteria combine biomineralization and subcellular compartmentalization to maintain intracellular iron reservoirs while limiting iron-dependent oxidative damage.

## Introduction

Iron is an essential micronutrient for nearly all living organisms, serving as a cofactor in numerous cellular processes, including respiration, DNA synthesis, and metabolism (1, 2). However, ferrous iron (Fe(II)) readily reacts with oxygen and reactive oxygen species (ROS) to generate highly toxic hydroxyl radicals through Fenton chemistry (3, 4). Iron can also compete with other transition metals, such as manganese and cobalt, for binding to metalloproteins, leading to enzyme mismetallation (1, 5). Cells must therefore balance iron acquisition with mechanisms that sequester excess iron while minimizing metal-dependent toxicity. In bacteria, iron storage is classically mediated by ferritin-like nanocages that oxidize and mineralize excess iron within a protein shell (6, 7). Recently, membrane-bounded iron-storage organelles termed ferrosomes were identified in several anaerobic bacteria, including both environmental and pathogenic species (8, 9). The gene cluster required for ferrosome biogenesis is conserved across diverse bacteria and archaea (9, 10), suggesting that ferrosomes represent a widespread strategy for intracellular iron compartmentalization.

Ferrosomes are lipid-bound compartments containing iron-rich biominerals. Recent studies suggest that ferrosomes promote adaptation to iron limitation under anaerobic conditions (9) and enable *Clostridioides difficile* to buffer transient iron spikes during host colonization (8). Despite their broad phylogenetic distribution and physiological importance, the chemical nature of ferrosome-stored iron remains poorly understood. Previous electron microscopy and elemental analyses demonstrated that ferrosomes are enriched in iron and phosphorus (8), but the oxidation state, coordination environment, and mineral organization of the stored iron have not been determined. Direct characterization of intracellular iron speciation within bacterial subcellular compartments remains technically challenging. X-ray absorption spectroscopy (XAS) has been widely used to characterize iron biominerals and metalloproteins (11, 12). In bacteria, XAS and complementary x-ray microscopy have been applied to characterize mixed-valence intracellular iron granules in *Shewanella putrefaciens* (13) and to investigate iron mineralization in magnetosomes of magnetotactic bacteria (14–17). These studies provide a foundation for investigating the chemical nature of ferrosome iron directly in intact cells.

The physiological consequences of storing large amounts of iron within ferrosome compartments also remain unclear. This question is particularly relevant in *C. difficile*, a leading cause of antibiotic-associated colitis and hospital-acquired infections in the United States (18). During infection, *C. difficile* encounters fluctuating iron availability and oxidative stress associated with host immune responses (2). Although a strict anaerobe, vegetative *C. difficile* cells can tolerate transient oxygen exposure and survive under low-oxygen conditions within the inflamed gastrointestinal tract (19). Because iron can participate in oxidative chemistry, understanding how ferrosomes compartmentalize iron and how this stored iron responds to oxygen exposure may provide insight into bacterial iron homeostasis and stress responses during infection.

Here, we investigated the chemical composition, redox properties, and physiological consequences of ferrosome-stored iron in *C. difficile*. To define the oxidation state and local coordination environment of the ferrosome biomineral, we applied Fe K-edge XAS and extended X-ray absorption fine structure (EXAFS) analysis to both isolated ferrosomes and intact cells, initially focusing on a *fur*::CT strain that overproduces these organelles. We further examined wild-type cells under conditions that promote ferrosome abundance, as well as ferrosome-deficient and complemented strains, to determine whether the iron speciation observed in the overproducing strain is also characteristic of ferrosomes under native genetic conditions. We also assessed the effects of oxygen exposure on ferrosome iron and investigated how ferrosome formation influences cellular responses to iron availability and oxidative stress. This integrated approach allowed us to examine the relationship between iron biomineralization, intracellular compartmentalization, and the redox behavior of ferrosome-stored iron.

## Results

### Characterization of the oxidation state of iron in isolated ferrosomes

To determine the redox state of ferrosome iron, we isolated ferrosomes from a *C. difficile fur*::CT mutant that constitutively overproduces these organelles (8). Scanning transmission electron microscopy paired with energy-dispersive X-ray spectroscopy (STEM-EDS) confirmed that the isolated ferrosomes are iron- and phosphate-rich biominerals (Fig. 1A-F), consistent with previous analyses showing an approximately 1:1 Fe:P ratio (8). We then collected Fe K-edge XAS spectra to determine the oxidation state and coordination environment of the ferrosome-stored iron.

**Fig. 1.**
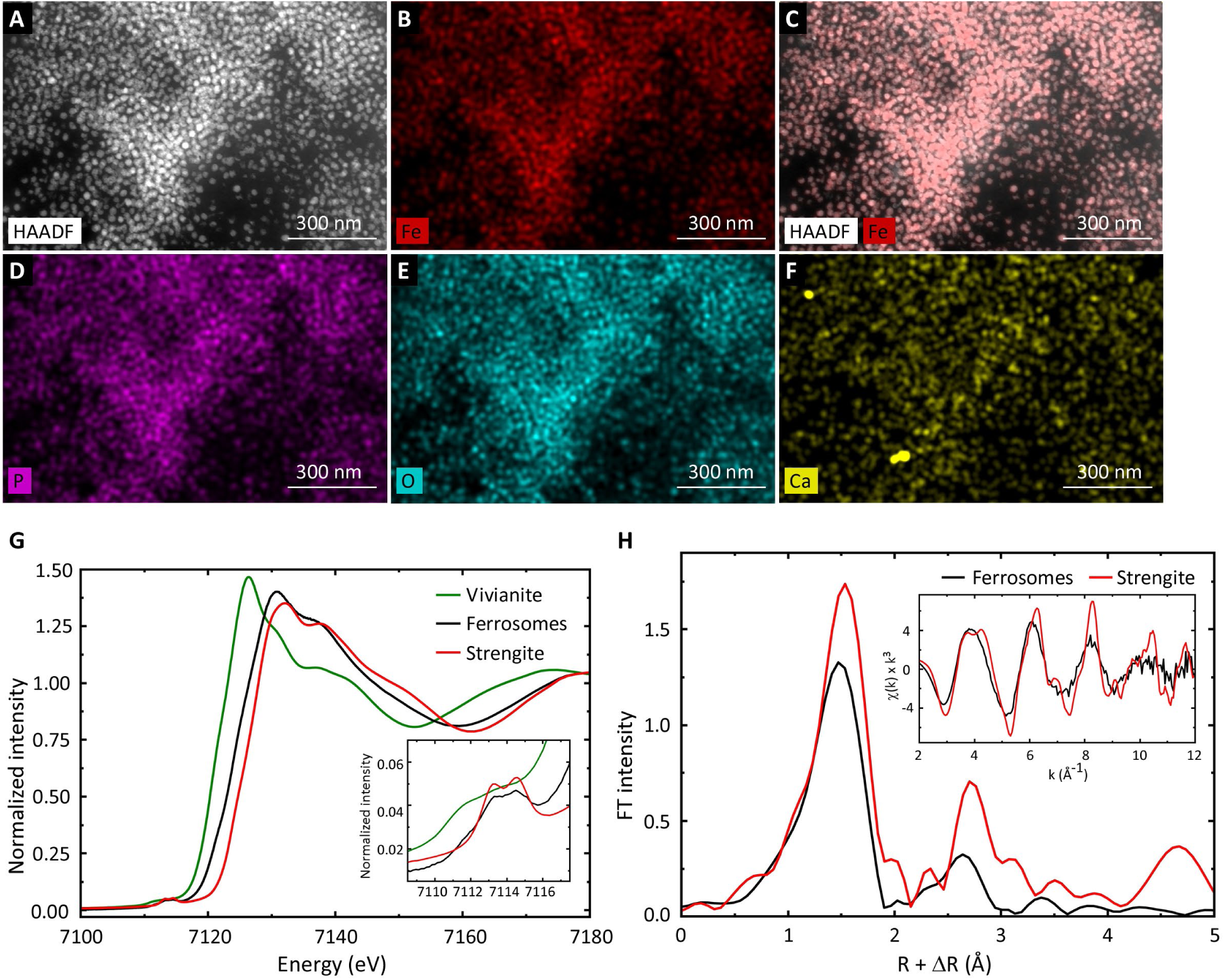
Isolated ferrosomes are mixed-valent Fe-phosphate minerals. (**A**) Representative HAADF-STEM micrographs of ferrosomes isolated from *C. difficile fur*::CT cells and corresponding EDS elemental maps showing (**B**) Fe, (**C**) merged HAADF with Fe signals, (**D**) P, (**E**) O, and (**F**) Ca. (**G**) Fe K-edge XAS spectra of isolated ferrosomes and mineral standards; the inset highlights the pre-edge region. (**H**) Fe K-edge EXAFS spectra (inset) and corresponding Fourier transforms (FT) of ferrosomes and strengite.

To interpret the ferrosome spectra, we compared them with representative ferric and ferrous phosphate mineral standards spanning a range of oxidation states and coordination environments (Fig. S1). The rising-edge position of the ferrosome spectrum is most similar to ferric phosphate phases, particularly strengite (FePO₄·2H₂O) (Fig. 1G). Higher-energy XANES features typical of hydrated ferric phosphates, arising from multiple scattering within the Fe–O–P framework, are also present but are shifted to lower energy and broadened, indicating structural disorder and the presence of Fe(II). Pre-edge analysis supported a mixed-valence assignment. Whereas strengite shows transitions consistent with high-spin Fe(III) in distorted octahedral coordination and vivianite (Fe_3_(PO_4_)_2_·8H_2_O) exhibits the three-feature pattern expected for Fe(II)O₆ coordination (Fig. S2, Table S3), the ferrosome pre-edge is dominated by Fe(III)-like features while retaining a lower-energy contribution characteristic of Fe(II). Linear-combination fitting using vivianite and strengite yielded an approximate composition of 34% Fe(II) and 66% Fe(III) (Table S4).

To define the local coordination environment, we collected EXAFS spectra of isolated ferrosomes and compared them with strengite (Fig. 1H). The ferrosome EXAFS is remarkably similar to strengite: the oscillations remain in phase, though their amplitude decays more rapidly at high k, indicative of increased structural disorder. The Fourier transform is dominated by a first-shell O peak and weaker second-shell features attributable to P at R’ = 2–3 Å (Fig. 1H). Quantitative fitting of the standards yielded average Fe–O distances of 1.99 Å for strengite and 2.08 Å for vivianite (Table S5), reflecting the smaller ionic radius of Fe(III) (4, 5). Strengite was best modeled using a split first shell comprising four shorter phosphate-derived Fe–O bonds and two longer water-derived Fe–O bonds. The ferrosome spectrum was best fit by the same 4+2 coordination model with an average Fe–O distance of 2.02 Å (Table S5), slightly longer than that of strengite but substantially shorter than that of vivianite. Together, these results identify the biomineral in isolated ferrosomes as a structurally disordered iron phosphate containing both Fe(II) and Fe(III), with iron remaining six-coordinated by phosphate- and water-derived oxygen ligands.

Notably, independently prepared ferrosome samples exhibited substantial variation in XAS spectra and inferred Fe(II)/Fe(III) ratios (Fig. S3 and Table S4). Because ferrosome isolation was performed aerobically, this variability strongly suggests that ferrosome iron undergoes oxidation during sample preparation. We therefore next examined ferrosomes directly in intact cells to better preserve their native redox state.

### Whole-cell XAS reveals the oxidation state of ferrosome iron in intact cells

Direct determination of the oxidation state of intracellular iron in intact bacterial cells by XAS is challenging because signals from specific subcellular compartments can be masked by iron distributed throughout the cell. In our previous study, WT *C*. *difficile* cells were grown under ferrosome-promoting conditions involving supplementation of BHIS medium with 150 µM of the iron chelator EDDHA, followed by addition of 200 µM FeSO_4_. These conditions yield only ∼10 ferrosomes per cell (8), an abundance insufficient for reliable in situ XAS analysis of ferrosome-associated iron. To overcome this limitation, we analyzed the ferrosome-overproducing *fur*::CT mutant grown in BHIS medium, in which ferrosomes frequently form large intracellular clusters (8) (Fig. 2B). Given the high iron content of these structures, ferrosome-associated iron was expected to make a substantial contribution to the bulk cellular Fe XAS signal relative to other intracellular iron pools. This strategy enabled direct in situ characterization of iron speciation in intact cells without organelle isolation. WT cells grown in standard BHIS medium do not produce detectable ferrosomes (Fig. 2A), and therefore served as a background control. The *fur::CT* Δ*fezB* mutant, which is defective in ferrosome biogenesis because of the deletion of *fezB* (8), served as an additional background control (Fig. 2C). Notably, energy dispersive spectroscopy (EDS) analysis of the irregular iron precipitates accumulated by the double mutant revealed that, unlike ferrosomes, these structures lacked characteristic iron and phosphorus enrichment and instead contained other metals, including zinc and magnesium (Fig. 2C and Fig. S4).

**Fig. 2.**
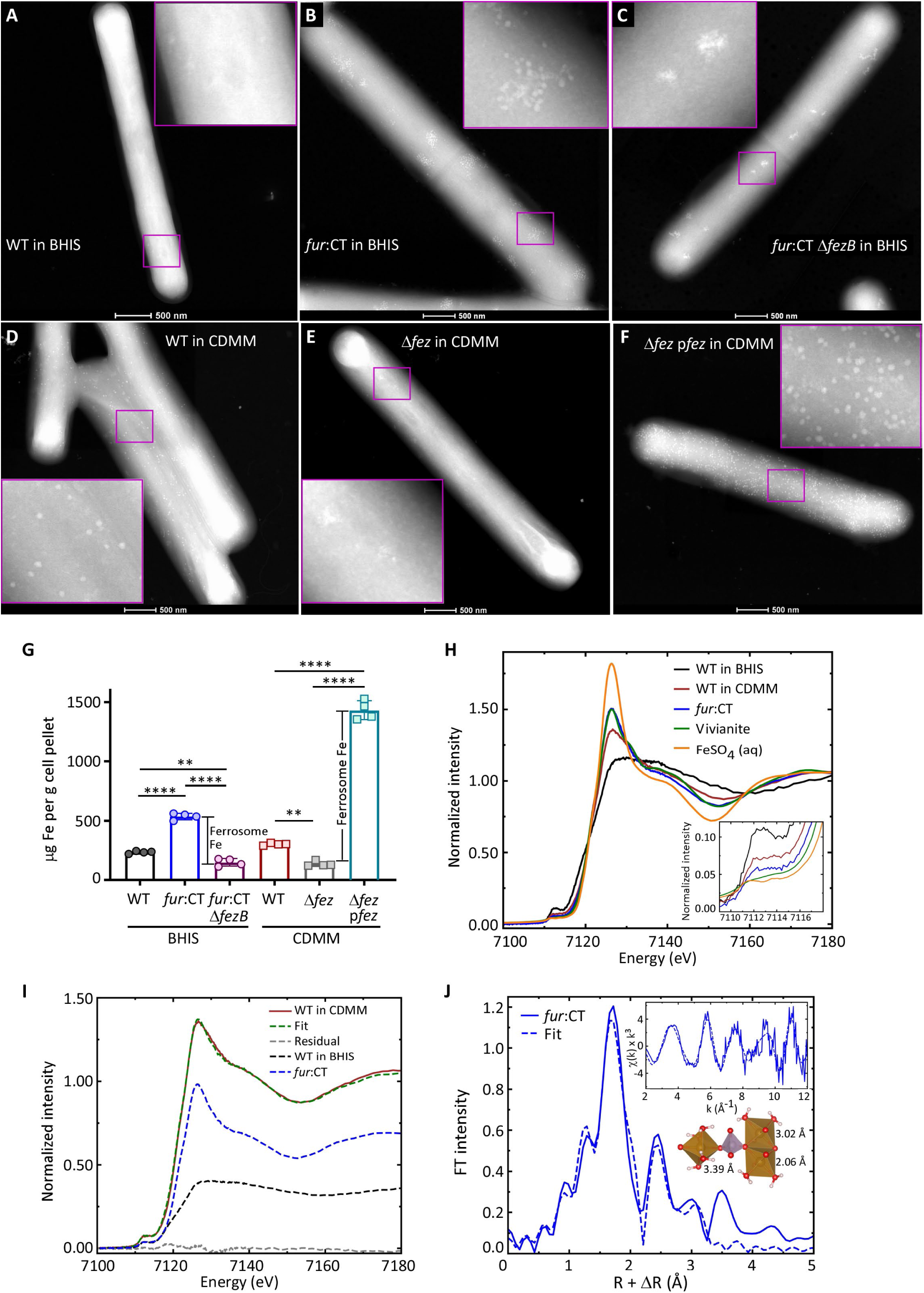
Characterization of ferrosomes in intact cells. (**A**-**C**) Representative STEM images of *C. difficile* WT (**A**), *fur*:: CT (**B**), and *fur*::CT Δ*fezB* (**C**) cells grown in BHIS medium, (**D**-**F**) Representative STEM images of WT (**D**), Δ*fez* (**E**), and Δ*fez*-p*fez* (**F**) cells grown under CDMM-based ferrosome-promoting conditions. (**G**) Intracellular iron levels in *C. difficile* WT, *fur*::CT and *fur*::CT Δ*fezB* strains grown in BHIS, and WT, Δ*fez*, and Δ*fez*-p*fez* strains grown in CDMM, as determined by ICP-MS. Data are expressed as µg of iron per gram of cell pellet. Experiments were performed independently twice. Data shown are one representative experiment (n=4). Statistical significance was determined by a one-way ANOVA: **, P<0.01; ****, P<0.0001. (**H**) Fe K-edge XAS spectra of WT and *fur*::CT cells grown in BHIS and WT cells grown under CDMM-based ferrosome-promoting conditions, compared with vivianite and aqueous ferrous sulfate reference standards. The inset shows the pre-edge region. (**I**) Linear combination fit of the Fe K-edge XAS of WT cells grown in CDMM, using the spectra of WT and *fur*::CT cells grown in BHIS as reference components. Dashed lines represent weighted reference spectra. (**J**) Fe K-edge EXAFS spectra (inset) and corresponding Fourier transform of *fur*::CT cells. The schematic illustrates the EXAFS fitting model based on the vivianite structure and the fitted bond distances.

To quantify the contribution of ferrosomes to the cellular iron pool, we measured intracellular iron levels by ICP-MS. The *fur::CT ΔfezB* mutant contained significantly less intracellular iron than WT cells (Fig. 2G), despite constitutive derepression of Fur-regulated iron uptake systems, suggesting activation of an iron-sparing or other compensatory iron homeostasis response in this background. In contrast, the *fur*::CT mutant accumulated approximately twice the intracellular iron content as WT cells and four times as much as the *fur*::CT Δ*fezB* mutant (Fig. 2G), suggesting that ferrosome-associated iron accounts for roughly three quarters of the total intracellular iron pool in *fur::CT* cells. WT and *fur::CT* strains were therefore subjected to XAS analysis, whereas the iron signal from the *fur::CT ΔfezB* mutant was insufficient for reliable measurements.

The Fe K-edge XAS spectrum of WT cells grown in standard BHIS medium, under conditions that do not support ferrosome formation, was compared with aqueous FeSO_4_ (Fig. 2H). The edge energy of WT closely matches that of FeSO_4_, indicating that most intracellular Fe remained reduced during sample preparation. However, the WT spectral line shape differs substantially from that of solvated Fe(II), whose sharp white line and 7140 eV shoulder is consistent with octahedral Fe-aquo complex. In contrast, the WT spectral features are broader and less structured, indicating that intracellular iron in WT cells exists in heterogeneous protein- and metabolite-associated coordination environments rather than as simple aqueous Fe(II). The WT spectrum also exhibits a broad Fe(II)-like pre-edge feature (Fig. 2H), further supporting the presence of multiple intracellular Fe coordination environments.

The spectrum of *fur*::CT cells differs markedly from that of WT cells, demonstrating that disruption of Fur-mediated regulation alters not only the amount of intracellular iron but also its speciation. Compared with WT, *fur*::CT shows a sharper white line and a shoulder near 7140 eV, indicating a more uniform octahedral Fe environment and consistent with Fe mineralization rather than heterogeneous soluble iron species (Fig. 2H). Although *fur*::CT may still contain a minor soluble iron fraction, the pronounced spectral changes indicate that this contribution is limited. Notably, the rising edge of *fur*::CT closely overlays that of vivianite (Fig. 2H), demonstrating that the ferrosome-encapsulated iron remains reduced in intact cells. The higher-energy features of *fur*::CT are broader and less resolved than those of crystalline vivianite (Fig. 2H), likely reflecting the structural disorder characteristic of biominerals (20). Because ferrous phosphate minerals display high variability in their XAS spectra (Fig. S1B), the close correspondence in edge energy and overall spectral line shape strongly indicates that ferrosomes store a ferrous phosphate mineral with a vivianite-like local structure.

EXAFS analysis further supports this assignment. The spectrum of aqueous FeSO_4_ is dominated by a single solvent-shell scattering, while the whole-cell samples exhibit contribution of multiple frequencies (Figs. S5 and 2J). The WT spectrum is out of phase with *fur*::CT below *k* = 5 Å^-1^, consistent with differences in the first shell coordination environments between soluble cellular iron and ferrosome-associated mineral. The *fur*::CT EXAFS is best fit by six oxygen ligands with an average Fe-O distance of 2.06 Å. The second shell is well fit by a Fe-Fe at 3.02 Å and two Fe-P at 3.39 Å (Fig. 2J). These structural parameters are consistent with vivianite (21). Together, the XAS and EXAFS data of Fe accumulated in *fur*::CT cells indicate a vivianite-like ferrous phosphate biomineral compartmentalized within ferrosomes.

To determine whether *fur* disruption alters ferrosome Fe speciation, we optimized growth conditions to promote ferrosome formation in WT *C. difficile*. Briefly, cells were passaged twice in chemically defined iron-free minimal medium (CDMM) before supplementation with 200 μM FeSO₄ to promote intracellular iron loading. Under these conditions, STEM imaging showed approximately 60 ferrosomes per cell (Fig. 2D). Compared with WT cells grown in standard BHIS, WT cells under this CDMM-based ferrosome-promoting regimen exhibit a sharper Fe K-edge XAS white line and a spectral profile resembling that of *fur*::CT cells and vivianite (Fig. 2H). The WT CDMM spectrum could be reproduced by a linear combination of the *fur*::CT (65%) and WT BHIS (35%) spectra (Fig. 2I), indicating that ferrosomes in WT cells grown in CDMM contain Fe species similar to those in *fur*::CT cells, but contribute a smaller fraction of the total cellular Fe XAS signal. The Fe K-edge EXAFS of WT cells grown in CDMM also overlaps with that of *fur*::CT cells (Fig. S5), further supporting similar local Fe coordination environments in ferrosomes from the two strains.

To test whether these spectral features depend on the *fez* operon, we examined two additional strains under the same CDMM-based ferrosome-promoting conditions as WT cells: the operon deletion mutant Δ*fez* and the complemented strain Δ*fez*-p*fez*, which carries the *fez* operon on a multicopy plasmid under the control of its native promoter. ICP-MS measurements showed that Δ*fez* cells contained lower Fe content than WT cells grown in CDMM, whereas Δ*fez*-p*fez* accumulated roughly five times as much Fe as WT cells (Fig. 2G). STEM imaging confirms that Δ*fez* lacks detectable ferrosomes, while Δ*fez*-p*fez* produces more than 300 ferrosomes per cell (Fig. 2E, F). The Fe K-edge XAS of Δ*fez* exhibits a broad, featureless white line resembling that of WT cells grown in standard BHIS (Fig. S6), consistent with Fe coordinated by heterogeneous cellular ligands, and its edge energy remains consistent with Fe(II), indicating that loss of the operon does not alter the oxidation state of background cellular Fe. In contrast, the spectrum of Δ*fez*-p*fez* resembles those of *fur*::CT cells and vivianite (Fig. S6). Together, these results show that the vivianite-like ferrous phosphate spectral signature depends on the *fez* operon and is restored upon complementation, supporting its assignment to ferrosomes.

### Ferrosome biominerals undergo oxidation from ferrous to ferric phosphate upon air exposure

The partial oxidation of isolated ferrosomes prompted us to test the redox-sensitivity of the native biomineral by exposing *fur*::CT cells to air for 3 h prior to XAS analysis. Air exposure shifted the Fe K rising-edge by ∼4 eV to higher energy, bringing it close to strengite (Fig. 3A), and produced more structured white line and pre-edge features characteristic of ferric phosphate. These spectral changes indicate that the ferrosome biomineral is readily oxidized within intact cells upon air exposure.

**Fig. 3.**
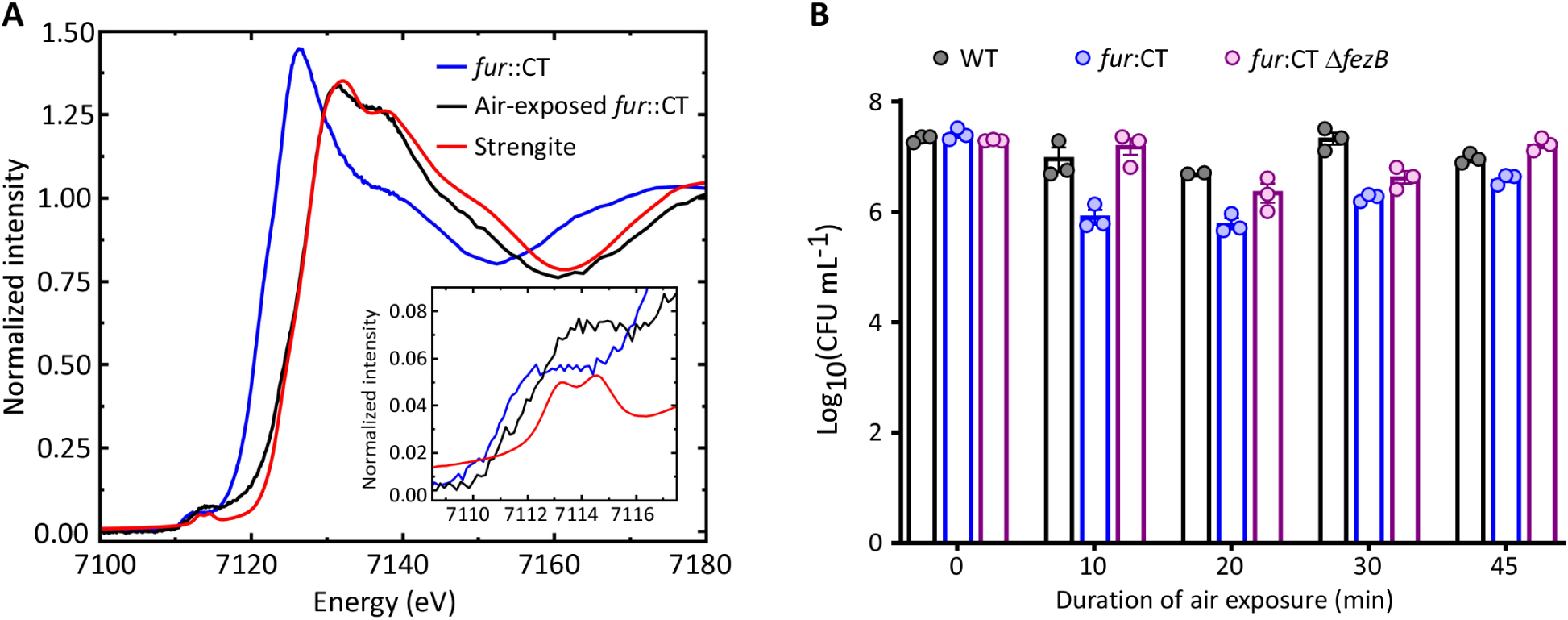
Ferrosome biominerals undergo oxidation upon air exposure. (A) Fe K-edge XAS spectra of *fur*::CT cells before and after air exposure, compared with strengite. (B) Survival of *C. difficile* WT, *fur*::CT and *fur*::CT Δ*fezB* strains following air exposure for the indicated times. Data are representative of one experiment with three biological replicates. The experiment was repeated independently four times with similar results. Statistical significance was determined using two-way ANOVA. No significant differences were observed between the strains at any given timepoint.

EXAFS analysis further supported this interpretation. Relative to anaerobically maintained *fur*::CT cells, air exposure contracts the first shell Fe-O distance by ∼0.05 Å (Table S5), consistent with oxidation. Notably, the oxidized whole-cell sample and isolated ferrosomes showed nearly identical average Fe-O distance (2.01 and 2.02 Å, respectively; Table S5), indicating that both undergo similar oxidation-dependent structural changes. Together, these data demonstrate that the ferrous phosphate biomineral compartmentalized within ferrosomes is highly redox-sensitive and readily converts to a ferric phosphate-like state upon oxygen exposure.

Although *C. difficile* is a strict anaerobe, vegetative cells can tolerate transient oxygen exposure and survive under low-oxygen conditions encountered within the host gastrointestinal tract (19, 22). We next tested whether ferrosome formation influences cellular survival during oxygen exposure. WT cells largely tolerated air exposure, exhibiting only ∼10% loss of viability after 45 min (Fig. 3B), consistent with previous observations (19, 22). The *fur*::CT Δ*fezB* mutant, which does not produce ferrosomes (Fig. 2C), exhibited air tolerance comparable to WT cells. In contrast, the *fur::CT* mutant, which constitutively produces numerous ferrosomes per cell (Fig. 2B), displayed a modest trend toward increased sensitivity to air exposure relative to the other two strains, however, this difference did not reach statistical significance, and the trend persisted throughout the 45 min exposure period (Fig. 3B). Together, these results suggest that although ferrosome-associated Fe(II) is readily oxidized upon oxygen exposure, ferrosome overproduction does not substantially compromise cellular oxygen tolerance under the conditions tested. Thus, bacterial cells can accommodate substantial intracellular stores of oxygen-sensitive ferrous phosphate without a major loss of oxygen tolerance.

### Ferrosome-associated iron has limited effects on intracellular iron bioavailability under iron-replete conditions

Many iron acquisition and homeostatic systems, including the ferrosome system, are constitutively derepressed in the *fur*::CT mutant (8, 23, 24). Consequently, constitutive expression of multiple iron acquisition systems may increase cellular iron demand. Consistent with this possibility, previous studies have shown that iron availability can remain limiting for optimal iron homeostasis even in nutrient-rich media (25–27). Given the substantial accumulation of redox-active Fe(II) within ferrosomes in the *fur*::CT mutant, we asked whether this stored iron contributes to the intracellular pool of bioavailable iron. To address this question, we used streptonigrin, an aminoquinone antibiotic that induces iron-dependent redox cycling and oxidative DNA damage (28–30). In several bacterial species, increased intracellular iron availability has been associated with greater streptonigrin sensitivity, making the antibiotic a potential indirect indicator of bioavailable iron (26, 30, 31). Surprisingly, despite the markedly elevated total cellular iron content in the *fur*::CT mutant (Fig. 2G), treatment with 4.9 μM streptonigrin caused only modest growth inhibition in all three strains under anaerobic conditions (Fig. 4A-B). Because streptonigrin activity depends in part on oxygen-dependent redox chemistry, which is likely limited under the anaerobic growth conditions (29), we supplemented cultures with a non-inhibitory concentration of H_2_O_2_ (75 μM) to enhance streptonigrin activity (Fig. 4C). Under these conditions, both *fur*::CT and *fur*::CT Δ*fezB* mutants exhibited greater streptonigrin sensitivity than WT cells. However, no significant difference was observed between the two mutants (Fig. 4D). Thus, although disruption of Fur-mediated iron homeostasis increased streptonigrin sensitivity, the substantially greater total iron accumulation in the ferrosome-overproducing *fur*::CT mutant did not result in greater sensitivity than in the ferrosome-deficient *fur*::CT Δ*fezB* mutant. These results suggest that ferrosome-associated iron does not proportionally increase the iron availability reflected by streptonigrin sensitivity.

**Fig. 4.**
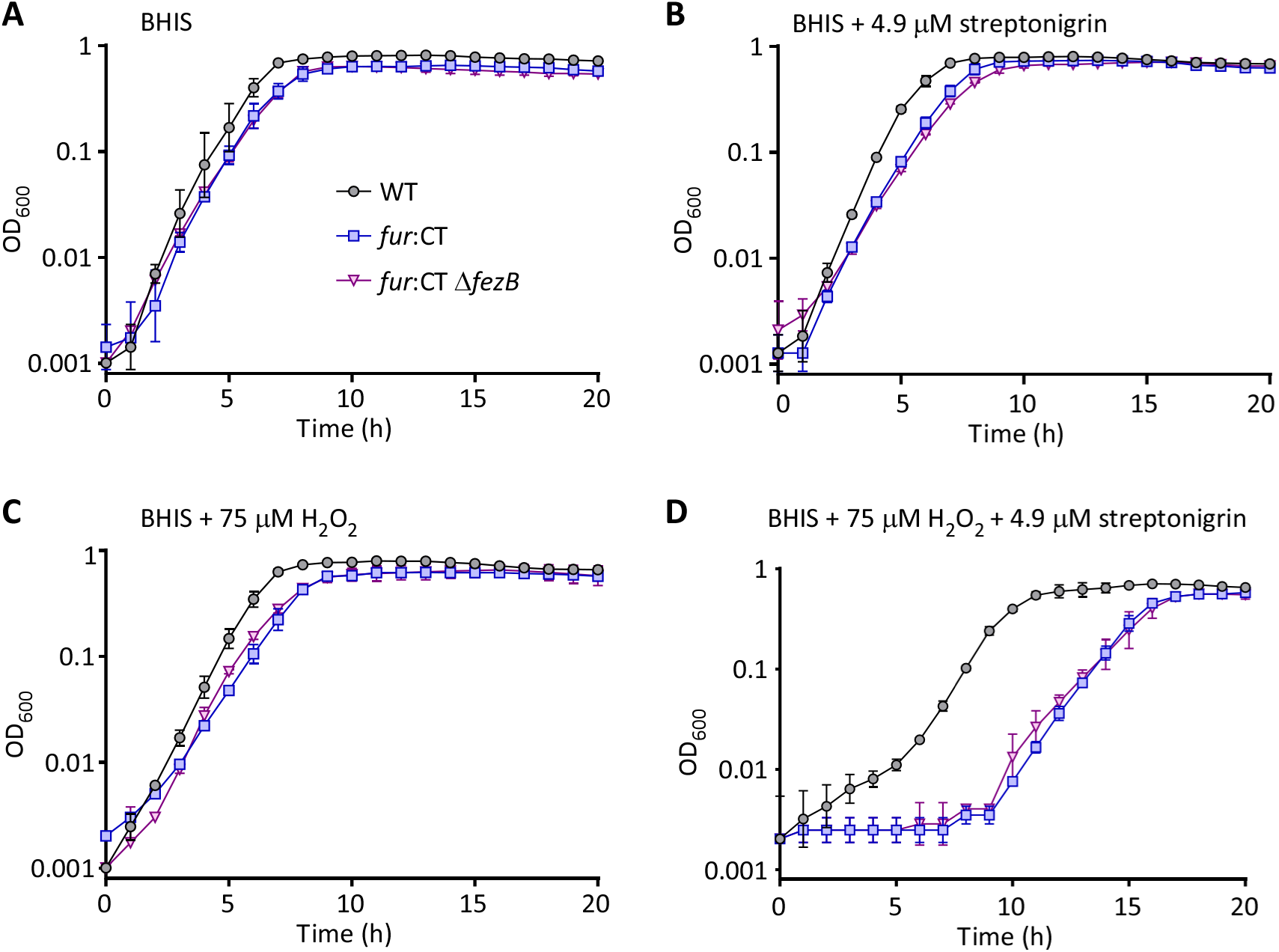
Ferrosomes have minimal impact on intracellular iron availability. Growth kinetics of *C. difficile* WT, *fur*::CT and *fur*::CT Δ*fezB* strains were monitored in the Coy anaerobic chamber under the following conditions: (**A**) BHIS, or BHIS supplemented with (**B**) 4.9 μM streptonigrin, (**C**) 75 μM H_2_O_2_, or (**D**) both 75 μM H_2_O_2_ and 4.9 μM streptonigrin. Experiments were performed three times independently, with three biological replicates per experiment. Data from one representative experiment are shown, with error bars indicating the standard deviation of the three biological replicates.

### Ferrosome-associated iron exhibits limited reactivity in oxidative reactions

During infection, *C. difficile* encounters reactive oxygen species such as hydrogen peroxide, generated by host immune cells (22, 32). To investigate whether ferrosome-associated iron influences cellular responses to oxidative stress, we examined the effects of H_2_O_2_ on WT, *fur*::CT, and *fur*::CT Δ*fezB* cells. In BHIS medium, all three strains exhibited similar growth kinetics (Fig. 5A). Supplementation with 100 or 150 μM H₂O₂ delayed the growth of both mutants relative to WT (Fig. 5B,C). These results indicate that disruption of Fur-mediated iron homeostasis increases susceptibility to peroxide-induced growth inhibition, but do not reveal a clear difference between the two mutants under these conditions. We next assessed cell viability following 15 min exposure to increasing H₂O₂ concentrations. Although survival was comparable among the three strains at lower concentrations, the *fur*::CT mutant exhibited significantly reduced survival at 180 μM H₂O₂ compared with both WT and the *fur*::CT Δ*fezB* mutant (Fig. 5D). Thus, deletion of *fezB* partially alleviated the increased peroxide sensitivity associated with Fur disruption. Consistent with activation of the peroxide stress response, H₂O₂ exposure increased expression of *rbr*, which encodes an H₂O₂-detoxifying enzyme, in all three strains (Fig. 5E).

**Fig. 5.**
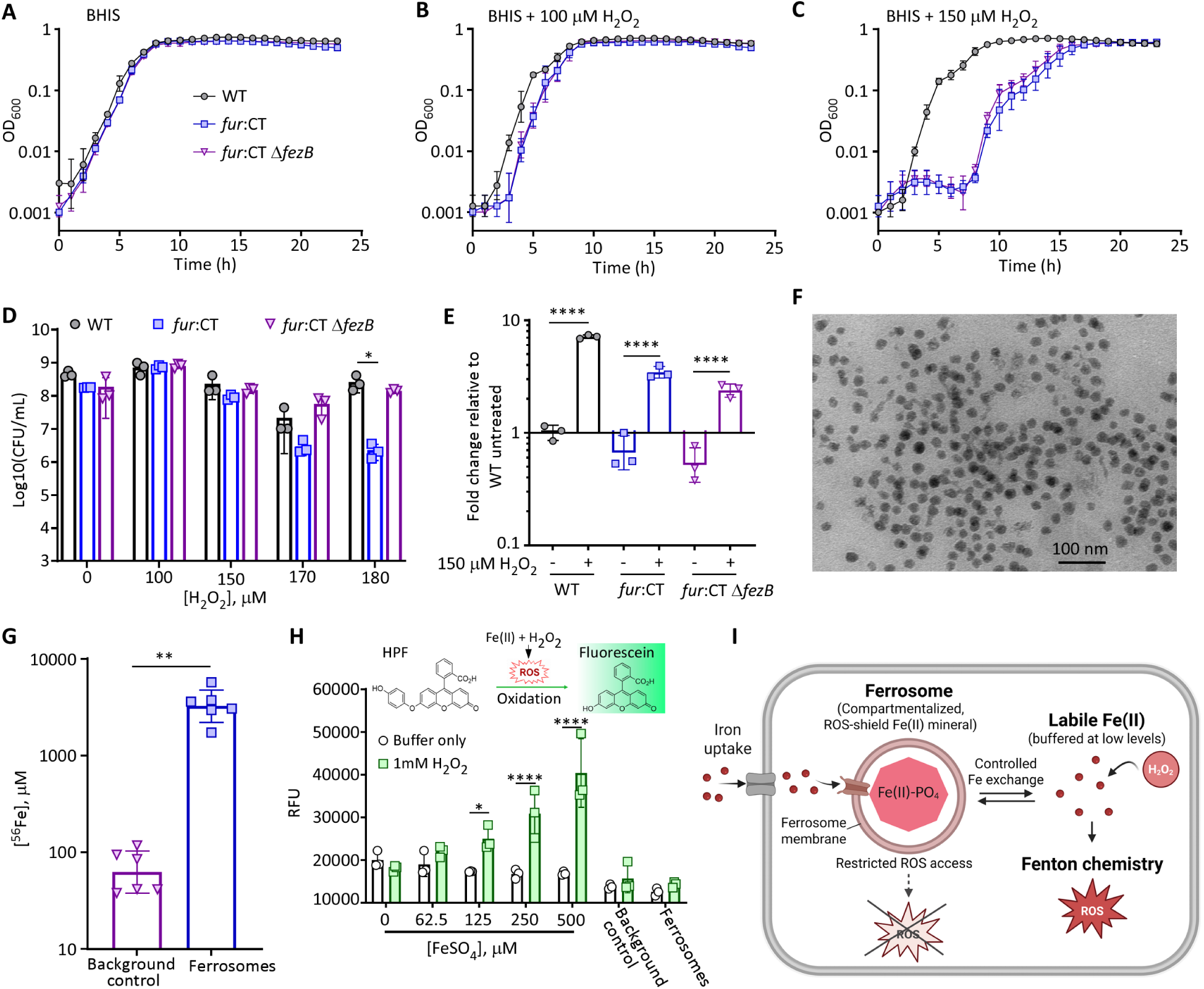
Ferrosomes sequester redox-active Fe(II) to limit oxidative stress. (**A**-**C**) Growth kinetics of *C. difficile* WT, *fur*::CT and *fur*::CT Δ*fezB* strains monitored in the Coy anaerobic chamber in BHIS (**A**) or BHIS supplemented with 100 µM (**B**) or 150 µM H_2_O_2_ (**C**). Experiments were performed independently three times, with three biological replicates per experiment. Data from one representative experiment are shown, with error bars indicating the standard deviation of the three biological replicates. (**D**) Viable cell counts (Log_10_CFU/mL) of the indicated strains following 15 min exposure to increasing concentrations of H_2_O_2_. Statistical significance was determined using two-way ANOVA: *P < 0.05. (**E**) Relative *rbr* transcript levels in the indicated strains following exposure to 150 μM H₂O₂ for 15 min, determined by qPCR. Transcript levels were normalized to the housekeeping gene *rpoB* and expressed relative to untreated WT cells. Statistical significance was determined using one-way ANOVA: ****P < 0.0001. (**F**) Representative transmission electron microscopy (TEM) image of ferrosomes isolated from *fur*::CT cells. (**G**) Iron concentration (μM) in ferrosome preparations isolated from *fur*::CT cells and the background control strain (*fur*::CT Δ*fezB*), as determined by ICP-MS. Data represent mean ± SD from two independent experiments, each with three biological replicates (n = 6). Statistical significance was determined using an unpaired Student’s *t*-test: **P < 0.01. (**H**) HPF fluorescence in reactions containing 1 mM H₂O₂ and increasing concentrations of FeSO₄ or ferrosomes isolated from *fur*::CT cells. The ferrosome-deficient *fur*::CT Δ*fezB* strain was used as a background control. Statistical significance was determined by a one-way ANOVA: *P < 0.05; ****P < 0.0001. (**I**) Proposed model illustrating the compartmentalization of Fe(II) within membrane-bound ferrosomes containing vivianite-like ferrous phosphate biominerals. The low solubility of ferrous phosphate minerals may limit the availability of soluble redox-active iron, while restricted access to mineral surfaces and limited diffusion of reactive intermediates across the ferrosome membrane may further constrain oxidative reactions.

To further examine whether iron sequestered within ferrosomes participates in oxidative reactions, we isolated ferrosomes from *fur*::CT cells (Fig. 5F) and quantified their iron content by ICP-MS. The isolated ferrosome fraction contained substantially more iron than the background control prepared from the ferrosome-deficient *fur*::CT Δ*fezB* strain (Figs. 5G and S7). We then monitored oxidative probe fluorescence using hydroxyphenyl fluorescein (HPF). HPF exhibits minimal fluorescence in its reduced state but becomes highly fluorescent upon oxidation by ROS, enabling sensitive detection of oxidative reactions driven by redox-active iron. As a positive control, FeSO_4_ induced a concentration-dependent increase in HPF fluorescence in the presence of H_2_O_2_ (Fig. 5H), demonstrating that the assay could detect iron-dependent fluorescein oxidation. In contrast, isolated ferrosomes produced no detectable increase in fluorescence despite containing approximately 4 mM iron (Fig. 5G-H). Although 50–60% of the ferrosome-associated iron can undergo oxidation during isolation (Table S4), an estimated 1.6–2 mM iron remains in the Fe(II) state. Because HPF readily penetrates lipid membranes, the negligible fluorescence signal observed with isolated ferrosomes suggests that ferrosome-stored Fe(II) has limited capacity to promote HPF oxidation under the conditions tested. Together, these findings support a model in which compartmentalization of Fe(II) within ferrous phosphate minerals limits its participation in oxidative reactions, while ferrosome-associated iron contributes to the increased peroxide sensitivity of *fur*::CT cells (Fig. 5I).

## Discussion

Our study demonstrates that ferrosomes are membrane-bounded compartments that sequester redox-active Fe(II) within bacterial cells. Using XAS and complementary elemental analyses, we show that ferrosomes in *C. difficile* predominantly contain a structurally disordered vivianite-like ferrous phosphate biomineral (Fig. 2H-J). Importantly, WT cells grown under conditions that promote ferrosome abundance exhibited similar XAS features, while deletion and complementation of the *fez* operon further supported the assignment of these features to ferrosomes. These results indicate that the Fe(II)-rich phosphate phase is not simply a consequence of altered iron homeostasis in the *fur*::CT strain. Early studies reported intracellular iron-rich inclusions in *Shewanella putrefaciens*, containing mixed-valence iron and phosphorus (13, 33), and Fe- and P-rich organelles in *Desulfovibrio magneticus* containing predominantly Fe(II) (17, 34). These observations, together with our findings in *C. difficile*, suggest that ferrous phosphate biomineralization represents a recurring iron-storage strategy among anaerobic bacteria. Unlike ferritin-like nanocages, which oxidize iron during storage, ferrosomes retain iron largely in the reduced Fe(II) state under anaerobic conditions (Fig. 2H-J). While ferritin stores ferric oxyhydroxide for iron buffering (35, 36) and magnetosomes biomineralize magnetite for magnetic navigation (37), ferrosomes accumulate a ferrous phosphate phase specialized for iron sequestration under reducing conditions. Together, these systems illustrate how iron mineral chemistry is tuned to distinct physiological demands and environmental conditions.

Our findings further suggest that membrane enclosure and phosphate mineralization enable cells to accumulate large intracellular Fe(II) stores while limiting the availability of soluble iron for cellular reactions. Although ferrosome-stored iron was oxygen-sensitive and became oxidized upon air exposure (Fig. 3), ferrosome-overproducing *fur*::CT cells did not exhibit substantially increased sensitivity to short-term air exposure (Fig. 3). The response to H₂O₂ was more complex. In growth kinetics experiments, both *fur*::CT and the ferrosome-deficient *fur*::CT Δ*fezB* mutant exhibited delayed growth at lower H₂O₂ concentrations relative to WT. However, the CFU survival assay revealed significant reduced survival of *fur*::CT cells at higher H₂O₂ concentrations compared with both WT and the Δ*fezB* mutant (Fig. 5). These results suggest that ferrosome formation contributes to the reduced survival of *fur*::CT cells under high-dose H₂O₂ stress, whereas the growth delay shared by both mutants may reflect broader changes in iron homeostasis resulting from constitutive Fur derepression.

Despite their substantially higher total cellular iron content (Fig. 2G), ferrosome-overproducing *fur*::CT cells showed streptonigrin susceptibility comparable to that of the ferrosome-deficient *fur*::CT Δ*fezB* mutant (Fig. 4). This observation suggests that the additional iron accumulated in ferrosomes does not substantially increase the iron availability detected by this assay. However, streptonigrin susceptibility has not been validated as a quantitative measure of the labile iron pool in *C. difficile*. Moreover, under the anaerobic, H₂O₂-supplemented conditions used here, streptonigrin activity may involve additional DNA-damaging mechanisms (28–30). Thus, these results provide indirect evidence that ferrosome-associated iron is relatively inaccessible to cellular reactions but do not establish the extent of iron exchange between ferrosomes and the cytosol.

The limited reactivity of ferrosome-associated iron was further supported by the observation that isolated ferrosomes did not produce a detectable increase in HPF fluorescence in the presence of H₂O₂ despite their high iron content (Fig. 5G-H). This result should be interpreted cautiously because ferrosome isolation and the HPF assay were performed outside the anaerobic chamber, potentially allowing partial oxidation of the mineral. Formation of an Fe(III)-phosphate-rich surface layer could passivate the mineral and restrict access to underlying Fe(II) (38), thereby reducing its reactivity. Moreover, the chemical specificity of HPF for individual ROS species has been questioned (39). Another potential explanation for the limited reactivity is the low solubility of ferrous phosphate minerals: under conditions of 1 mM total phosphate and pH 7, less than 1 µM Fe(II) remains dissolved in equilibrium with vivianite (Table S6). This low solubility would limit the pool of soluble iron available for redox reactions, potentially reducing ROS generation. Additional factors, including limited accessibility of mineral surfaces and restricted diffusion of reactive intermediates across the ferrosome boundary, may further contribute to the observed inhibition of oxidative chemistry. Nevertheless, oxidation of ferrosome-associated Fe(II) by O₂ could potentially generate superoxide, which may be converted to H₂O₂ and diffuse beyond the ferrosome compartment. The extent of ROS production during ferrosome oxidation, its contribution to cellular oxidative stress, and the extent to which cellular ROS-detoxification systems mitigate these effects remain unclear.

The ferrous phosphate composition of ferrosomes likely reflects adaptation to anaerobic physiology. Ferritin-mediated storage depends on iron oxidation (40), a strategy favored in aerobic environments where ferric minerals remain stable. In contrast, ferrosome gene clusters are primarily found in organisms adapted to anaerobic reducing environments in which Fe(II) predominates (8–10). Under such conditions, confinement of Fe(II) and phosphate within the organelle may favor precipitation of ferrous phosphate minerals. Fe(II) imported by the P_1B6_-type ATPase FezB may accumulate together with phosphate to drive mineral nucleation, while the reducing intracellular environment stabilizes the ferrous phase. These observations suggest that ferrosomes create a specialized compartment optimized for anaerobic iron sequestration.

Although the local iron coordination environment resembles vivianite, several observations suggest that the ferrosome biomineral exists in a more disordered or amorphous state. Ferrosomes contain a lower Fe:P ratio than crystalline vivianite and lack evidence of extensive long-range structural order despite prolonged intracellular residence (Fig. 2). Amorphous biominerals are increasingly recognized as kinetically favorable mineral states because they precipitate rapidly and possess lower kinetic barriers to formation than crystalline phases (41, 42). Such properties may be particularly beneficial in *C. difficile*, where ferrosomes appear to buffer transient increases in intracellular iron (8). The organelle membrane may additionally stabilize this metastable phase by restricting crystal growth and maintaining a confined chemical environment.

These findings are relevant to the physiology of *C. difficile* during infection, where the bacterium encounters fluctuating iron availability, oxidative stress, and intermittent oxygen exposure within the inflamed gastrointestinal tract (22, 32, 43). Our previous work demonstrated that ferrosomes promote adaptation to transient iron overload during colonization (8). This study further suggests that ferrosomes enable sequestration of excess Fe(II) as a phosphate mineral, potentially limiting its availability for cellular redox reactions, allowing *C. difficile* to maintain iron balance under constantly changing host-associated environments. The extent to which this storage strategy influences oxidative stress resistance and bacterial survival during infection remains to be determined.

Our work also highlights the utility of whole-cell XAS for characterizing intracellular metal speciation in intact bacterial cells. Previous studies have applied whole-cell XAS and spatially resolved x-ray methods to study magnetosome formation and iron speciation in intracellular iron-rich granules. These studies have demonstrated the ability of whole-cell EXAFS to characterize poorly ordered intracellular iron minerals, including phosphate-rich ferric hydroxide intermediates proposed to precede magnetite formation (15–17, 44–48). Because whole-cell XAS captures contributions from both ferrosomes and other cellular iron pools, we compared spectra of WT, *fur*::CT, Δ*fez*, and Δ*fez*-p*fez* strains to distinguish ferrosome-associated spectral features from background cellular iron without isolating the organelles. Indeed, ferrosome isolation resulted in partial oxidation of stored iron and substantial variability in iron speciation (Figs. 1, S3 and Table S4), underscoring the sensitivity of the biomineral to oxygen exposure during sample preparation. By analyzing intact cells, whole-cell XAS and EXAFS circumvented these isolation-associated artifacts and, together with STEM-EDS and ICP-MS, identified an amorphous ferrous phosphate phase with vivianite-like local coordination in situ (Figs. 1, 2). Although whole-cell XAS does not resolve the subcellular distribution of iron, spatially resolved approaches such as scanning transmission X-ray microscopy (STXM) could provide complementary information on the distribution and speciation of iron in ferrosomes and other intracellular pools. Combining these approaches may further advance the characterization of intracellular biominerals and metal-containing compartments under near-native conditions.

More broadly, our study expands current models of bacterial metal homeostasis by demonstrating that intracellular compartmentalization can mitigate toxicity associated with reactive metals. The widespread conservation of the ferrosome gene cluster across many obligate and facultative anaerobic bacteria and archaea suggests that this mechanism may be broadly important in anaerobic microbial physiology (9, 10). Future studies will be needed to determine how iron is mobilized from ferrosomes, how biomineral formation is regulated, and whether ferrosomes perform conserved or species-specific functions across diverse microorganisms.

Collectively, our findings establish ferrosomes as a distinct class of bacterial iron-storage organelles that sequester redox-active Fe(II) as a ferrous phosphate biomineral. These results reveal an alternative strategy for intracellular iron management and underscore the importance of subcellular compartmentalization in bacterial metal homeostasis.

## Methods

### Bacterial strains and growth conditions

*E. coli* strains used for cloning were grown aerobically in LB media supplemented with 25 µg/ml chloramphenicol and 10 µg/ml tetracycline, as appropriate. *C. difficile* strains were grown anaerobically at 37°C in an anerobic chamber (Coy Laboratory Products) maintained under an atmosphere of 90% nitrogen, 5% hydrogen, and 5% carbon dioxide. Unless otherwise indicated, *C. difficile* cultures were grown in brain heart infusion broth (RPI) supplemented with 0.5% yeast extract (BHIS). To promote ferrosome formation in WT C*. difficile*, cells were grown in iron-free CDMM (10 mg/ml casamino acids, 0.5 mg/ml L-tryptophan, 0.5 mg/ml L-cysteine, 5 mg/ml Na_2_HPO_4_, 5 mg/ml NaHCO_3_, 0.9 mg/ml KH_2_PO_4_, 0.9 mg/ml NaCl, 10 mg/ml glucose, 0.04 mg/ml (NH_4_)_2_SO_4_, 0.0196 mg/ml CaCl_2_, 0.02 mg/ml MgCl_2_ · 6H_2_O, 0.01 mg/ml MnCl_2_ · 4H_2_O, 0.001 mg/ml CoCl_2_ · 6H_2_O, 0.001 mg/ml D-biotin, 0.001 mg/ml calcium-D-pantothenate, 0.001 mg/ml pyridoxine). The *fur*::CT and *fur*::CT Δ*fezB* strains were cultured in BHIS supplemented with 20 μg mL^-1^ lincomycin.

### Plasmid Construction

All oligonucleotides used in this study are listed in Table S1, and all plasmids are listed in Table S2. To construct pKM197-*fezAB*-*fezA*g2 for deletion of *fezA* and *fezB* from the *C. difficile* genome, the region upstream of *fezA* was amplified with the primers “*fezA*-upstream_fwd” and “*fezA*-up-rev-doublemut” and the region downstream of *fezB* was amplified with the primers “*fezB*-down-fwd-doublemut” and “*fezB* downstream_rev”. These fragments were cloned into pKM197 digested with NotI and XhoI with Gibson assembly. The resulting plasmid was then digested with KpnI and MluI, and the sgRNA targeting *fezA, “fezA*_gRNA2_gBlock” was cloned into the plasmid using Gibson assembly.

To construct pKM197e-*fez*-*fezX*g3 to delete *fezX* from the Δ*fezA* Δ*fezB* strain to construct Δ*fezXAB*, the pKM197 plasmid was modified to allow for conjugation through *E. coli*. pKM197 was modified to construct pKM197e by replacing the Tn916 oriT for *Bacillus subtilis* conjugation with the TraJ oriT for *E. coli* conjugation. TraJ was amplified from pMSR0 (addgene plasmid # 161984) with the primer pair “E.coli_traJ_fwd”/”E.coli_traJ_rev”. The PCR product was digested with ApaI (NEB) and ligated with ApaI-digested pKM197 with T4 DNA ligase. A point mutation identified in *xylR* in pKM197e was also repaired by amplifying *xylR* from pCM111 with “*clsA*-*xylR*-UP-F” and “*clsA*-*xylR*-down-R”. The PCR product was digested with PacI and BglII and ligated into pKM197e digested with the same enzymes with T4 DNA ligase.

To construct pKM197e-*fezXAB*-*fezX*g3, the region downstream of *fezB* was amplified with the primers “fezB downstream_rev” and “fezB-down-fwd-fezdel”. The region upstream of *fezX* was amplified with “fezX upstream_fwd” and “fezX-up-fwd-fezdel”. These fragments were assembled into pKM197e digested with XhoI and NotI was Gibson assembly. This plasmid was then digested with KpnI and MluI to insert “fezX-gRNA3” using Gibson assembly.

To construct pDSW1728-*fezXAB*, the entire *fez* operon was amplified with the primers “fezX-pDSW1728-fwd-nhei” and “fezB-rev-pDSW1728-bamhi”. The PCR product was digested with NheI and BamHI and ligated into pDSW1728 digested with the same enzymes using T4 DNA ligase.

### *C. difficile* strain construction

To construct the Δ*fezA* Δ*fezB* strain, pKM197-*fezAB*-*fezA*g2 was first transformed into *E. coli* MG1655 to generate multimers of the plasmids. Plasmid DNA extracted from MG1655 was then transformed into *B. subtilis* JH BS2. The plasmids were transferred from *B. subtilis* JH BS2 to *C. difficile* CD196 through conjugation. Briefly, 100 µl of *B. subtilis* grown in BHI (37 g/L BHI) supplemented with 2.5 µg/ml tetracycline and 2.5 µg/ml chloramphenicol and 100 µl of *C. difficile* grown in BHIS. The mixture was spotted onto a BHIS agar plate and incubated overnight at 37°C under anaerobic conditions. The mating plate was scraped, and the cells were resuspended in BHIS and plated onto BHIS agar plates supplemented with 50 µg/ml kanamycin and 10 µg/ml thiamphenicol. Transconjugants were patched onto BHIS plates supplemented with 10 µg/ml tetracycline and BHIS plates supplemented with 10 µg/ml thiamphenicol. Transconjugants which were thiamphenicol-resistant and tetracycline-sensitive were plated on TY agar (3% Tryptone, 2% yeast extract, 0.1% sodium thioglycolate) supplemented with 1% xylose and 10 µg/ml thiamphenicol for three passages. Deletion of *fezA* and *fezB* was screened by PCR using the primers “fezX-merodiploid-conf-fwd” and “fezB-conf-rev-3”. The pKM197 plasmid was cured from the strains by passaging ten times in BHIS.

To construct the Δ*fezXAB* strain, pKM197e-*fezXAB*-*fezX*g3 was transformed into *E. coli* HB101 and transferred by conjugation into Δ*fezA* Δ*fezB* strain. Briefly, HB101 *E. coli* harboring the plasmid was grown on LB agar plates supplemented with 25 µg/mL chloramphenicol and 10 µg/mL tetracycline. Prior to mating, 400 µl of mid-log *C. difficile* culture was heat-shocked at 50°C for 10 minutes. The HB101 *E. coli* donor strain was resuspended into the heat-shocked *C. difficile* culture. The mixture was spotted onto BHI agar plates and incubated at 37°C for 16 hours. BHI plates were scraped and transconjugants were selected on BHIS agar plates supplemented with 50 µg/ml colistin and 10 µg/ml thiamphenicol. Transconjugants were plated onto TY agar supplemented with 1% xylose and 10 µg/ml thiamphenicol. Mutants were screened by PCR using the primers “fezX_ck2_fwd” and “fezB_ck2_rev”. The pKM197e plasmid was cured by passaging the strain ten times in BHIS. Gene deletions were confirmed by PCR, Sanger sequencing, and whole genome sequencing. To construct the Δ*fez* p*fez* complementation strain, pDSW1728-*fezXAB* was introduced into Δ*fezXAB* through conjugation using *E. coli* HB101 as the donor.

### Ferrosome isolation

*C. difficile fur*::CT cells were grown overnight in BHIS broth supplemented with 20 μg/mL lincomycin at 37°C and subcultured at a 1:50 dilution into 1 L of fresh BHIS medium supplemented with 20 μg/mL lincomycin. Cultures were grown for 8 hours. Cells were harvested and resuspended in lysis buffer (10 mM Tris-HCl, pH 8.0, 50 mM NaCl, 1 mM EDTA, and 250 mM sucrose) supplemented with EDTA-free cOmplete protease inhibitor cocktail (Roche). Cells were lysed by sonication, and cell debris was removed by centrifugation at 12,000 rpm for 30 min at 4°C. Clarified lysates were layered onto a 65% sucrose cushion and centrifuged at 35,000 rpm for 2 h at 4°C. The resulting pellet was washed with 50 mM Tris-HCl (pH 8.0), resuspended in 50 μL of the same buffer, loaded into XAS cuvettes, flash-frozen in liquid nitrogen, and stored at −80°C prior to analysis.

### XAS sample preparation

For XAS measurements of strains grown in BHIS, *fur*::CT, WT, and *fur::CT* Δ*fezB* strains were grown overnight in BHIS broth at 37°C, diluted 1:50 into fresh BHIS, and grown for an additional 6 h at 37°C. For XAS measurements of *C. difficile* strains grown in CDMM, WT, Δ*fez*, and Δ*fez* + pDSW1728-*fezXAB* (p*fez*) were grown overnight in iron-free CDMM at 37°C. Cultures were diluted 1:50 into fresh iron-free CDMM and grown for 24 hours at 37°C. 200 µM FeSO_4_ was then added to the cultures and incubated for 3 hours before cell harvesting. Following growth, aliquots (10 mL) of culture were removed from the anaerobic chamber and centrifuged at 4,000 rpm for 7 min at room temperature. Cell pellets were immediately returned to the Coy anaerobic chamber and resuspended in 0.5× PBS that had been equilibrated in the chamber overnight or longer to deplete residual oxygen. Cells were washed three times by centrifugation and resuspension in anaerobically equilibrated 0.5× PBS, with all washing steps performed inside the chamber to minimize oxygen exposure. Following the final wash, the OD_600_ was measured, and cells were resuspended to an OD_600_ of 10 in 0.5× PBS. Samples were then loaded into cuvettes sealed with 25 µm-thick Kapton tape, flash frozen in liquid nitrogen, and shipped overnight on dry ice for XAS analysis.

### H_2_O_2_ CFU Assay

*C. difficile* strains were grown overnight in BHIS at 37°C, subcultured 1:50 into fresh BHIS, and grown to mid-log phase at 37°C. Cultures were adjusted to an OD₆₀₀ of 0.2, followed by the addition of H₂O₂ and incubation at 37°C for 15 min. Following treatment, cultures were serially diluted, and 2.5 μL aliquots of each dilution were spotted onto BHIS agar plates to determine colony-forming units (CFUs).

### RT-qPCR

*C. difficile* strains were grown overnight in BHIS at 37°C, subcultured 1:50 into fresh BHIS, and grown to mid-log phase at 37°C. Cultures were either left untreated or treated with 150 μM H₂O₂ for 15 min. Cells were harvested by centrifugation, and cell pellets were stored at −80°C until RNA extraction. Cells were lysed using Lysing Matrix B tubes (MP Biomedicals, catalog no. 6911100) and a FastPrep-24 bead beater. Total RNA was extracted using the RNeasy Mini Kit (Qiagen) according to the manufacturer’s instructions, with each sample eluted in 60 μL of RNase-free water. RNA samples were treated with TURBO DNA-free DNase (Ambion) and quantified using a NanoDrop spectrophotometer. For cDNA synthesis, 200 ng of total RNA from each sample was used with the High-Capacity cDNA Reverse Transcription Kit (Applied Biosystems). Quantitative PCR was performed using iQ SYBR Green Supermix (Bio-Rad) on a CFX96 real-time PCR system (Bio-Rad). The housekeeping gene *rpoB* gene was used as an internal reference for normalization.

### X-ray absorption spectroscopy

Upon receipt, cell suspensions or isolated ferrosomes were stored in liquid nitrogen until measurement. A natural strengite sample (Indian Mountain, Alabama, USA) was procured and thoroughly ground with cellulose and placed between Kapton film for measurement. XAS spectra were collected at the Stanford Synchrotron Radiation Lightsource on beamline 7-3. A Si(220) double crystal monochromator, oriented at Φ=0, was used for energy selection at Fe K-edge. The harmonic rejection was provided by detuning the monochromator to 60%. Scattering background was reduced using a Mn filter placed in front of Soller slits. During data collection, samples were maintained at ∼10 K under He atmosphere using a Cryo Industries closed-loop He-cryocooler. Fe K-edge XAS of cell pellets and isolated ferrosome were collected in fluorescence mode using a Canberra multi-element, solid-state, Ge detector. Strengite spectrum was collected in transmission mode using ionization chambers. To minimize beam damage, four spots per sample cuvette were analyzed with two to eight scans collected at each spot. No evidence of beam damage was observed. For all measurements, an Fe foil reference spectrum was simultaneously collected in transmission mode for energy calibration. The vivianite spectrum was measured using the same protocol at room temperature in transmission mode, using a sample from Stanford Mineral Collection.

For data processing, detector channels exhibiting high background signals from ice diffraction were excluded, and the remaining channels were averaged in ATHENA (49). Spectra were normalized and energy calibrated by setting the Fe foil edge energy, defined as the first maximum in first derivative, to 7112 eV. Normalized spectra were imported into PySpline (50) for post-edge background subtraction. EXAFS was extracted by setting E_0_ to 7130 eV and fitting the background using a 3-region spline (with polynomial orders 2, 3, and 3). Fitting of *k*^3^-weighted EXAFS was performed using EXAFSPAK (51), and theoretical scattering amplitudes and phase functions were generated with FEFF7 (52). Initial model geometries were adopted from reported mineral structures. Linear combination analysis was performed in ATHENA.

### Inductively coupled plasma mass spectrometry (ICP-MS)

*C. difficile* strains were grown under the same conditions used for XAS experiments. For whole-cell metal quantification, aliquots (10 mL) of culture were collected in metal-free tubes, pelleted, and washed three times with 1× PBS. The mass of each cell pellet was recorded prior to acid digestion. Pellets were resuspended in 300 μL of 70% nitric acid and 75 μL of 30% trace metal-grade hydrogen peroxide and incubated overnight at 60°C. Samples were then diluted with 11.7 mL of ultrapure water and further diluted 10-fold in 1.75% nitric acid before ICP-MS analysis.

For ICP-MS analysis of isolated ferrosomes, ferrosomes were isolated from 1 L of *fur*::CT culture. The ferrosome-deficient *fur*::CT *ΔfezB* was processed in parallel as a negative control. Isolated ferrosomes were resuspended in 200 μl of wash buffer (100 mM HEPES pH 7.0, 0.85% NaCl, 100 mM sucrose). A 10 μL aliquot of the isolated ferrosome suspension was subjected to acid digestion using the same procedure described above. Before ICP-MS analysis, digested samples were diluted 20-fold in 1.75% nitric acid.

Metal-free tubes and pipette tips were used throughout sample preparation to minimize metal contamination. Elemental concentrations were determined using a NexION 5000 ICP-MS (PerkinElmer) at the Yale Analytical and Stable Isotope Center. For whole-cell samples, ^56^Fe measurements were corrected for sample dilution and normalized to the corresponding cell pellet mass.

### Scanning transmission electron microscopy (STEM) and transmission electron microscopy (TEM)

For cellular STEM imaging, *C. difficile* strains were grown in the same conditions used for XAS measurements. Cells (5 mL) were harvested by centrifugation and washed three times with 0.1 M HEPES buffer (pH 7.0). Washed cells were deposited onto Formvar/carbon-coated copper grids (EMS, cat# FCF400-Cu-50) and allowed to absorb for 25 min. Grids were then fixed with 2.5% glutaraldehyde in 0.1 M HEPES buffer for 30 min, followed by three washes with water. STEM imaging was acquired using an FEI Tecnai Osiris transmission electron microscope operated at 200 kV in high-angle annular dark-field scanning transmission electron microscopy (HAADF-STEM) mode at the Yale Institute for Nanoscience and Quantum Engineering.

For TEM imaging of isolated ferrosomes, ferrosomes were resuspended in 200 μL of buffer containing 100 mM HEPES, 0.85% NaCl, and 100 mM glycerol. The suspension was serially diluted twice, and 8 μL of the diluted sample was deposited onto a glow-discharged (30 s) carbon-coated 200-mesh copper grid. After incubation for 1 min, excess liquid was removed by blotting with filter paper. Grids were air-dried at room temperature for 10 min prior to imaging. Electron micrographs were acquired using a JEOL JEM-1400Plus transmission electron microscope operated at 80 kV.

### Air Shock assay

*C. difficile* strains were grown overnight in BHIS broth at 37°C and subcultured 1:50 into fresh BHIS medium. Cultures were grown to mid-log phase at 37°C, and the OD_600_ of each culture was adjusted to 0.2 in BHIS medium. Normalized cultures were serially diluted in 1× PBS, and 2.5 μL of each dilution was spotted onto BHIS agar plates that had been equilibrated in the Coy anaerobic chamber overnight or longer to deplete residual oxygen. Plates were removed from the anaerobic chamber and exposed to ambient air in a biosafety cabinet with the lids removed for 10, 20, 30, or 45 min. Following air exposure, plates were returned to the anaerobic chamber and incubated overnight at 37°C prior to colony enumeration.

### Growth kinetic assays

For all growth assays, 96-well plates were equilibrated in the Coy anaerobic chamber for at least 72 hours prior to use to deplete residual oxygen. *C. difficile* strains were grown overnight in BHIS at 37°C, then subcultured 1:50 into fresh BHIS and grown to mid-log phase at 37°C. Cultures were adjusted to an OD_600_ of 0.1 and diluted 1:100 into 200 µl of growth medium in the equilibrated 96-well plates. For streptonigrin assays, a 0.5 mg/mL streptonigrin (Sigma) stock solution was prepared in DMSO and diluted 1:200 into BHIS with or without 75 µM H_2_O_2_. For H_2_O_2_ sensitivity assays, cells were inoculated into BHIS containing 100 µM or 150 µM H_2_O_2_. Plates were incubated at 37°C in a Biotek plate reader (Agilent) located inside the Coy anaerobic chamber, with continuous shaking, and OD_600_ measurements were recorded every 15 min.

### Hydroxyphenyl fluorescein (HPF) assay

Hydroxyphenyl fluorescein (HPF; MedChemExpress) was used as a fluorescent probe to assess oxidative reactions associated with isolated ferrosomes. HPF exhibits minimal fluorescence in its reduced form but becomes fluorescent upon oxidation by highly reactive oxygen species, including hydroxyl radicals and other oxidants. This assay was used to determine whether iron contained within isolated ferrosomes promotes oxidative reactions in the presence of H_2_O_2_. Ferrosomes were isolated from 1 L cultures of the *fur*::CT strain and resuspend in 200 µl of wash buffer (100mM HEPES pH 7.0, 0.85% NaCl, 100mM sucrose). The *fur*::CT Δ*fezB* strain, which does not produce ferrosomes, was processed in parallel and used as a background control. HPF was prepared as a 0.2 mM stock solution in N,N-Dimethylformamide (DMF), and 1 µl was added to each well of a black-walled, clear-bottom 96-well plate to achieve a final concentration of 1 µM in a total reaction volume of 200 µl. For FeSO₄ control reactions, 2 μL of a 50 mM FeSO₄ stock solution prepared in 0.1 M HCl was added to wells containing wash buffer, with or without H₂O₂. For ferrosome reactions, 50 μL of isolated ferrosome suspension or the corresponding background control preparation was added to wells in the presence or absence of 1 mM H₂O₂. Because ferrosome isolation was performed outside the Coy anaerobic chamber, the HPF assay was also conducted under ambient atmospheric conditions. Fluorescence was measured immediately after the addition of FeSO₄ or ferrosome preparations using an Agilent BioTek plate reader (excitation, 487 nm; emission, 528 nm).

## Data Availability

All relevant data are within the manuscript and its Supplementary Information.

## Acknowledgments

We thank members of the Pi Laboratory for critical comments of the manuscript. This work was supported by the National Institutes of Health grant DP2 AI184552 (H.P.) and the Searle Scholars Program through a Searle Scholar Award (SSP-2025-109, H.P.). Use of the Stanford Synchrotron Radiation Lightsource, SLAC National Accelerator Laboratory, is supported by the U.S. Department of Energy, Office of Science, Office of Basic Energy Sciences under Contract No. DE-AC02-76SF00515. The SSRL Structural Molecular Biology Program is supported by the DOE Office of Biological and Environmental Research, and by the National Institutes of Health, National Institute of General Medical Sciences (P30GM133894). The contents of this publication are solely the responsibility of the authors and do not necessarily represent the official views of NIGMS or NIH.

## Author Contributions

K.M.F. and H.P. conceived and designed the experiments. K.Z. performed the XAS and EXAFS experiments, Y.L. isolated ferrosomes and performed the TEM imaging experiments, M.J.A provided experimental support during the EXAFS measurements and procured the strengite and vivianite datasets, K.M.F. performed all other experiments. K.Z., K.M.F., R.S., and H.P. wrote the paper with input from all authors. All authors reviewed the paper.

