## Supplementary material for "Ferrosome Organelles Compartmentalize a Redox-Active Ferrous Phosphate Biomineral": SI

[*]These authors contributed equally.

**Table S1. Oligonucleotides and gene fragments used in this study.**

| Name | Sequence |
|---|---|
| FezA-upstream-fwd | caggaaacagctatgaccgcgcggccgctaaatatcaagaactttctaaaatac |
| fezA-up-rev-doublemut | atataatggggatttacattatcctttcatttagttagattttg |
| fezB-down-fwd-doublemut | ataatgtaaatccccattatataaaacatatgtatataaaaaatttataattc |
| fezB downstream_rev | cttcttatttttatgctagcctcgagattgatttgaagg |
| E.coli_traJ_fwd | gcgaggaagcggaagagcgcccaatacgcagggccccctgcttcggggtca |
| E coli traJ rev | aatttatctacaattttttatcctgcagggggcccgatcggtcttgccttg |
| clsA-xylR-UP-F | caaaattcccttcaaatcaatctcgaggctagcataaaaataagaagcctgcatttg |
| clsA-xylR-down-R | cacttttacctctatttttatcagatcttgttaatactttattatctatag |
| fezB-down-fwd-fezdel | atgctatacttaatttccccattatataaaacatatgtatataaaaaatttataattc |
| fezX upstream_fwd | caggaaacagctatgaccgcgcggccgcaactaaattgttgtttttattc |
| fezX-up-fwd-fezdel | ttttatataatggggaaattaagtatagcatgtgtaaatggtattttc |
| fezX-pDSW1728-fwd-nhei | gtaagctagccatttataataatatatctaacgcactttaat |
| fezB-rev-pDSW1728-bamhi | tcagggatccttaatagtagttataagtattggtgttttccaat |
| fezX-merodiploid-conf-fwd | actctgggtcactcgttatac |
| fezB-conf-rev-3 | atatgtaaccatgtatgcaaatatcc |
| fezX_ck2_fwd | gtgtatataaattccattctttcc |
| qPCR_rpoB_fwd | TGCTGTTGAAATGGTTCCTG |
| qPCR_rpoB_fwd | cggttggcatcatcattttc |
| rbr_qPCR_F | TGCAGGAGAGTCAGAAGCAA |
| rbr_qPCR_R | GCATGTTCCTTTTCATTGTTAGC |
| fezA_gRNA2_gBlock | taattaaactgtaaaggtacggtaccggaagacttatctacatctagttttagagctagaaatagcaagttaaaataaggctagtccgttatcaacttgaaaaagtggcaccgagtcggtgctttttttctatggagaaatctagatcagcatgatgtctgactagacgcgtttaagctctgcaactatttt |
| fezX_guide3 | taattaaactgtaaaggtacggtaccgttactgtattttaaagttttgttttagagctagaaatagcaagttaaaataaggctagtccgttatcaacttgaaaaagtggcaccgagtcggtgctttttttctatggagaaatctagatcagcatgatgtctgactagacgcgttaagctctgcaactattttt |

**Table S2. Plasmids and strains used in this study**

| **Plasmids** | **Description** | | **Reference** |
|---|---|---|---|
| pKM197 | Addgene plasmid #132665 | | (1) |
| pCM111 | Encodes WT *xylR* and *cas9* | | (2) |
| pDSW1728 | Addgene plasmid #120812. Complementation plasmid | | (3) |
| pMSR0 | addgene plasmid #161984. Encodes oriT TraJ | | (4) |
| pKM197-fezAB-fezAg2 | Used to delete *fezA* and *fezB* from *C. difficile* | | This study |
| pKM197e-fezXAB-fezXg3 | Used to construct *fezXAB* deletion mutant in *C. difficile* | | This study |
| pKM197e | pKM197 plasmid modified to allow for conjugation through E. coli and a repaired *xylR* gene | | This study |
| pDSW1728-fezXAB | For expression of entire fez operon in *C. difficile*, under the control of its native promoter | | This study |
| **Species** | **Genotype** | **Description** | **Source** |
| *C. difficile* CD196 | WT | Wildtype *C. difficile* strain | Lab stock |
| *C. difficile* CD196 | Δ*fez* | Deletion of *fezXAB* operon | This study |
| *C. difficile* CD196 | Δ*fez +* pDSW1728-*fezXAB* | Δ*fezXAB* strain complemented with the entire *fez* operon | This study |
| *C. difficile* CD196 | *fur::CT* | Clostron insertion mutation of *fur* | Lab stock |
| *C. difficile* CD196 | *fur::CT* Δ*fezB* | Clostron insertion mutation of *fur* and deletion of *fezB* | (5) |
| *Escherichia coli* DH5a | | | Lab stock |
| *E. coli* MG1655 | | | Lab stock |
| *E. coli* HB101 | pRK24 | For conjugation of plasmids into C. difficile | Lab stock |
| *Bacillus subtilis* 168 | JH BS2 | Harbors Tn916 for conjugation of plasmids into C. difficile | Lab stock |

**Table S3. Fe K-pre-edge fitting results.**

| Sample | σ/γ | Voigt 1 | | Voigt 2 | | Voigt 3 | |
|---|---|---|---|---|---|---|---|
| | | Center | Amplitude | Center | Amplitude | Center | Amplitude |
| Isolated ferrosome | 0.526 | 7111.7 | 0.0180 | 7113.2 | 0.0512 | 7114.6 | 0.0446 |
| Strengite | 0.392 | | | 7113.2 | 0.0434 | 7114.6 | 0.0424 |
| Vivianite | 0.714 | 7111.2 | 0.0252 | 7111.9 | 0.0240 | 7113.5 | 0.0384 |

**Table S4. Linear combination fitting of isolated ferrosome.**

| Sample | %Vivianite | %Strengite |
|---|---|---|
| Isolated ferrosome 1 | 34 | 66 |
| Isolated ferrosome 2 | 48 | 52 |

**Table S5. Fe K-edge EXAFS curve-fitting results.**

| Sample | Coordination/path | $R$ (Å)[a] | $\sigma^2$ (Å$^2$×10$^5$) | $\Delta E_0$ (eV) | $k$-range | $F^b$ |
|---|---|---|---|---|---|---|
| *fur*::CT | 2 Fe-O | 1.93 | 765 | -9.23 | 2-12 | 0.484 |
| | 4 Fe-O | 2.12 | 295 | | | |
| | 1 Fe-Fe | 3.02 | 446 | | | |
| | 2 Fe-P | 3.39 | 547 | | | |
| Vivianite | 6 Fe-O | 2.08 | 1394 | -7.85 | 2-12 | 0.307 |
| | 1 Fe-Fe | 3.07 | 1068 | | | |
| | 2 Fe-P | 3.15 | 1770 | | | |
| Isolated ferrosome | 4 Fe-O | 1.97 | 514 | -5.24 | 2-12 | 0.233 |
| | 2 Fe-O | 2.11 | 381 | | | |
| | 4 Fe-P | 3.25 | 1644 | | | |
| Strengite | 4 Fe-O | 1.97 | 545 | -2.96 | 2-12 | 0.281 |
| | 2 Fe-O | 2.04 | 485 | | | |
| | 4 Fe-P | 3.29 | 657 | | | |
| $FeSO_4$ (aq) | 6 Fe-O | 2.11 | 393 | -8.30 | 2-12 | 0.295 |
| *fur*::CT oxidized | 6 Fe-O | 2.01 | 650 | -3.03 | 2-10 | 0.409 |

[a]The estimated standard deviations for the bond lengths are on the order of ±0.02 Å.

[b]The error is given by $[\Sigma k^6(\chi_{exptl}-\chi_{calcd})^2/\Sigma k^6\chi_{exptl}^2]^{1/2}$. The $S_0^2$ factor was set at 1.

**Table S6.** Calculation of $Fe^{2+}$ concentration in equilibrium with vivianite solid at pH 7, 25 $^{O}$C. The calculation was performed using the Geochemist's Workbench 2026 (www.gwb.com).

| Species | $Fe^{2+}$ | $PO_4^{3-}$ | $Na^+$ | Vivianite |
|---|---|---|---|---|
| Input | 0 | 1 mM total ($H_3PO_4$+$H_2PO_4^-$+$HPO_4^{2-}$+$PO_4^{3-}$) | 1.5 mM | ∞ |
| Equilibrium | 0.85 µM | 0.0026 µM | 1.5 mM | ∞ |

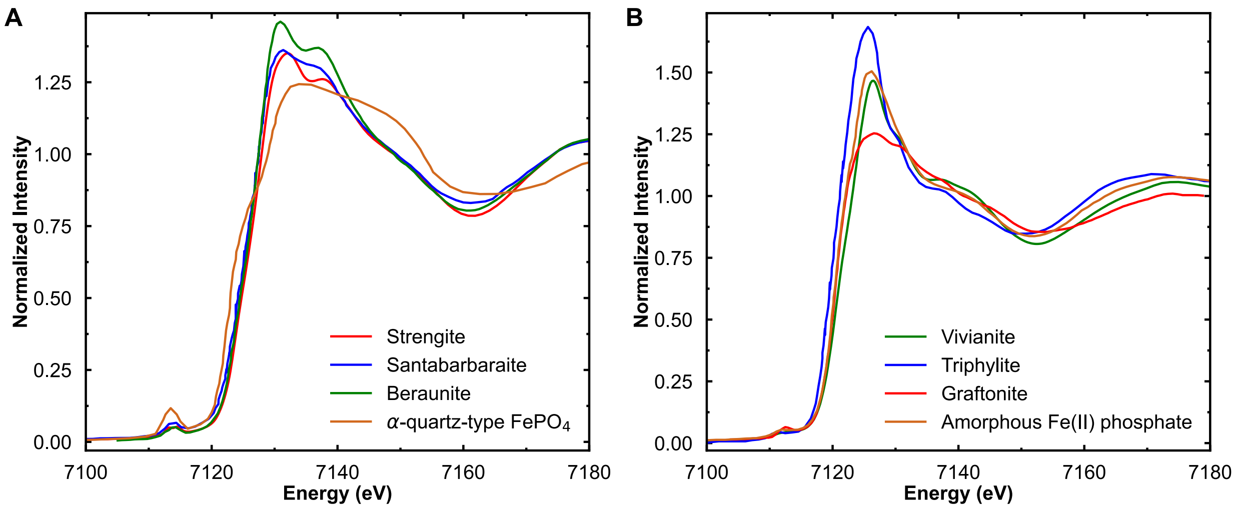


**Figure S1.** Fe K-edge XAS spectra of reference iron phosphate minerals. (A) Ferric phosphates: strengite ($FePO_4 \cdot 2H_2O$), santabarbaraite ($Fe_3(PO_4)_2(OH)_3 \cdot 5H_2O$) (6), beraunite ($Fe^{2+}Fe^{3+}{}_5(PO_4)_4(OH)_5 \cdot 4H_2O$) (7), and α-quartz-type $FePO_4$ (rodolicoite) (8). (B) Ferrous phosphates: vivianite ($Fe_3(PO_4)_2 \cdot 8H_2O$), triphylite ($LiFePO_4$) (9), graftonite ($(Fe,Mn,Ca)_3(PO_4)_2$) (10), amorphous ferrous phosphate (11). Spectral data were digitized from published figures using WebPlotDigitizer (12).

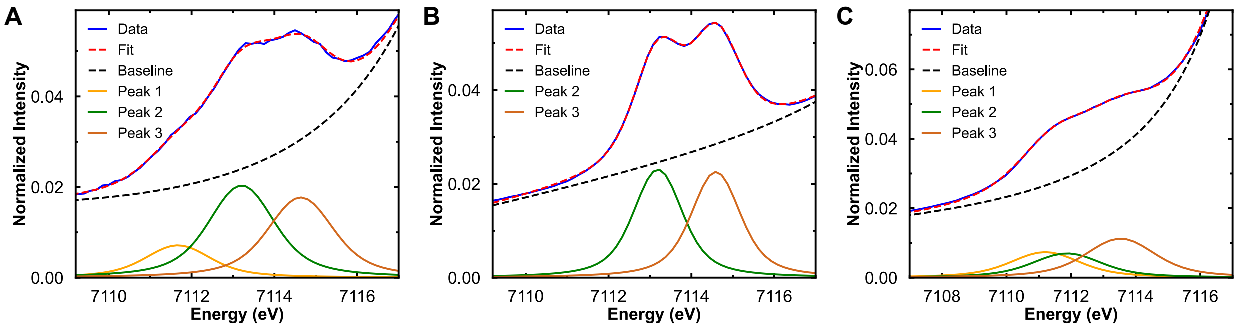


**Figure S2.** Fe K-pre-edge fits of (A) isolated ferrosomes, (B) strengite, and (C) vivianite.

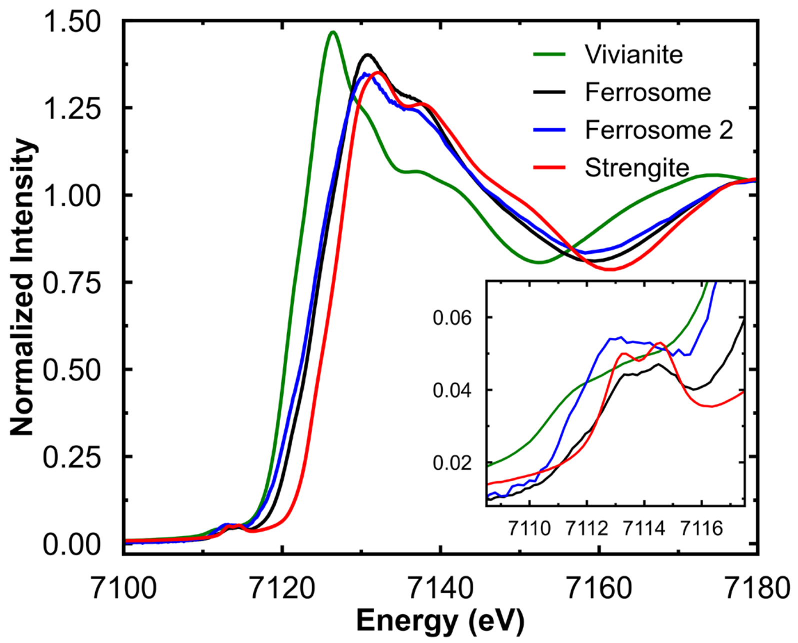


**Figure S3.** Fe K-edge XAS of isolated ferrosome from independent preparation batches. The inset shows an expanded view of the pre-edge region.

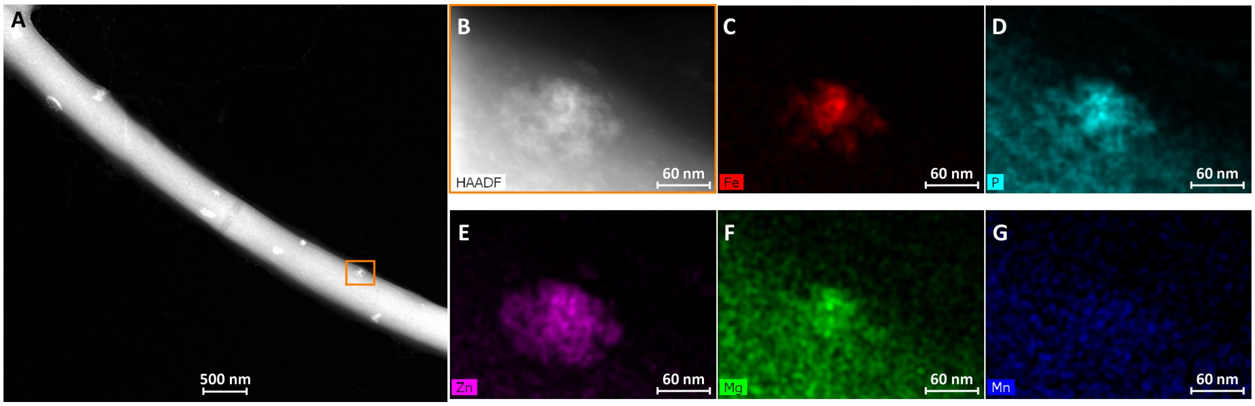


**Figure S4.** Electron-dense, metal-containing granules in the *fur*::CT *fezB* mutant visualized by STEM-EDS. (A–B) Representative high-angle annular dark-field scanning transmission electron microscopy (HAADF-STEM) images of *fur*::CT Δ*fezB* cells and corresponding energy-dispersive X-ray spectroscopy (EDS) elemental maps showing the distribution of (C) Fe, (D) P, (E) Zn, (F) Mg, and (G) Mn. Unlike ferrosomes, these granules do not exhibit characteristic Fe/P enrichment but instead contain other metals, including Zn and Mg.

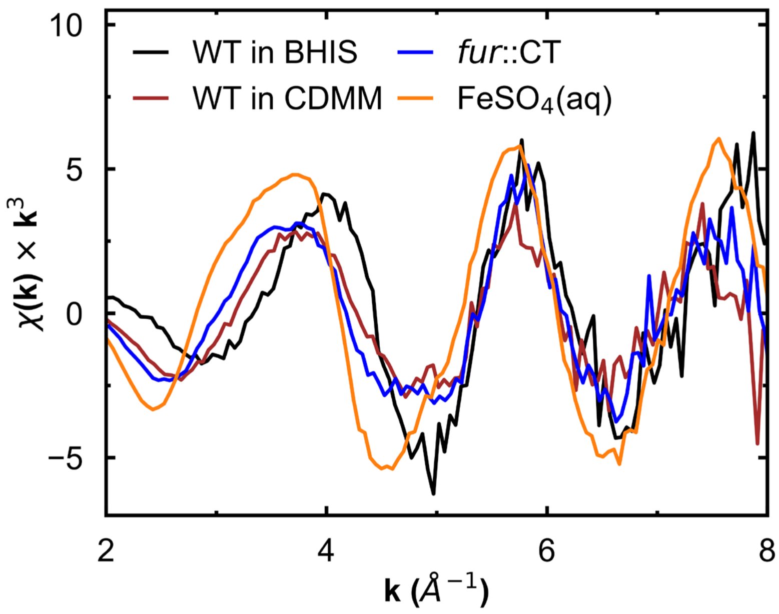


**Figure S5.** Fe K-edge EXAFS spectra of WT cells grown in BHIS (non-ferrosome-promoting conditions) or CDMM (ferrosome-promoting conditions), *fur*::CT cells grown in BHIS (constitutive ferrosome formation), and an aqueous $FeSO_4$ standard (aq).

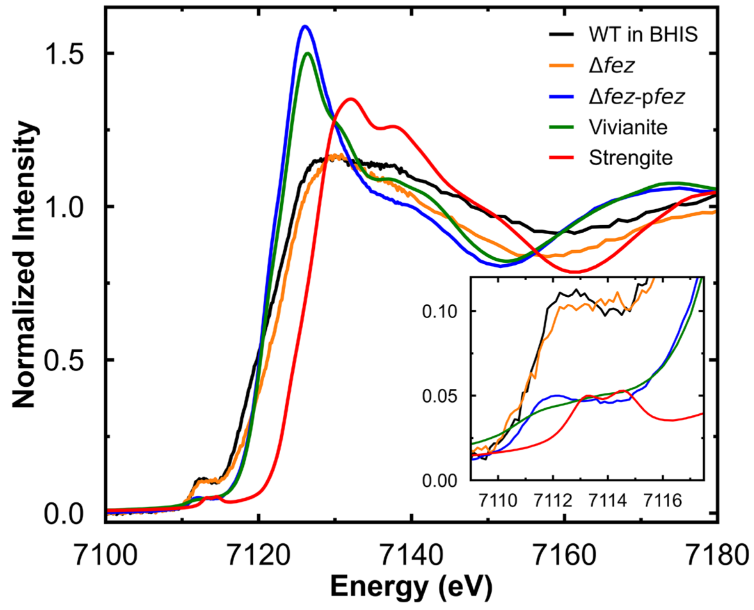


**Figure S6.** Fe K-edge XAS spectra of the Δ*fez* deletion strain and the operon-complemented strain (Δ*fez*-p*fez*) grown in CDMM (ferrosome-promoting conditions), compared with WT cells grown in BHIS (non-ferrosome-promoting conditions) and reference iron-phosphate minerals.

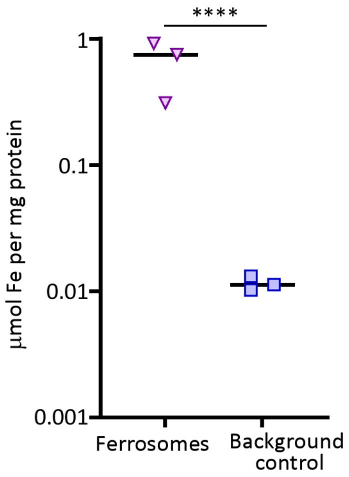


**Fig. S7.** Iron enrichment in ferrosome-containing fractions relative to recovered protein. Iron content in ferrosome fractions isolated from *fur*::CT cells and the background control strain (*fur*::CT Δ*fezB*) was quantified by ICP-MS. Iron content was normalized to the protein concentration of the final purified fractions, measured using a BCA assay, and expressed as µmol Fe per mg protein. Data represent mean ± SD from three biological replicates. Statistical significance was determined using Welch's *t*-test: ****P < 0.0001.
